# Cpk2, a catalytic subunit of cyclic AMP-PKA, regulates growth and pathogenesis in rice blast

**DOI:** 10.1101/173104

**Authors:** Poonguzhali Selvaraj, Qing Shen, Fan Yang, Naweed I. Naqvi

## Abstract

The cAMP-Protein Kinase A signalling, anchored on CpkA, is necessary for appressorium development and host penetration, but indispensable for infectious growth in *Magnaporthe oryzae*. In this study, we identified and characterized the gene encoding the second catalytic subunit, *CPK2*, whose expression was found to be lower compared to *CPKA* at various stages of pathogenic growth in *M. oryzae*. Deletion of *CPK*2 caused no alterations in vegetative growth, conidiation, appressorium formation, or pathogenicity. Surprisingly, the *cpkA*Δ*cpk2*Δ double deletion strain displayed significant reduction in growth rate and conidiation compared to the single deletion mutants. Interestingly, loss of *CPKA* and *CPK2* resulted in morphogenetic defects in germ tubes (with curled/wavy and serpentine growth pattern) on hydrophobic surfaces, and a complete failure to produce appressoria therein, thus suggesting an important role for *CPK2*-mediated cAMP-PKA in surface sensing and response pathway. *CPKA* promoter-driven *CPK2* expression partially suppressed the defects in host penetration and pathogenicity in the *cpkA*Δ. Such ectopic *CPK2* expressing strain successfully penetrated the rice leaves, but was unable to produce proper secondary invasive hyphae, thus underscoring the importance of CpkA in growth and differentiation *in planta*. The Cpk2-GFP localized to the nucleus and cytoplasmic vesicles in conidia and the germ tubes. The Cpk2-GFP colocalized with CpkA-mCherry on vesicles in the cytosol, but such overlap was not evident in the nucleus. Our studies indicate that CpkA and Cpk2 share overlapping functions, but also play distinct roles during pathogenesis-associated signalling and morphogenesis in the rice blast fungus.

## Introduction

The Protein kinase A (PKA) family of Ser/Thr kinases is highly conserved in eukaryotes, and serves important phosphorylation-dependent functions in signal transduction and development (Hanks & Hunter, 1995). The PKA holoenzyme is an inactive heterotetramer composed of two regulatory (R) and two catalytic (C) subunits and the cooperative binding of two cAMP molecules to the R subunit frees and activates its C subunits to phosphorylate hundreds of targets and regulate a vast swath of metabolism and cellular physiology.

The catalytic subunit of PKA (PKA-C) is a typical structure for protein kinases and PKA signaling plays a central role in vegetative growth, development, mating, stress response, and pathogenicity in various fungi (D’Souza & Heitman, 2001, Lengeler *et al.*, 2000). Multiple PKA isoforms are expressed in mammalian cells and have tissue-specific roles indicative of functional diversity. Three *TPK* genes encoding PKA-C were identified in the budding yeast *Saccharomyces cerevisiae*, and subsequently shown to share redundant and distinct functions in viability and in pseudohyphal morphogenesis, respectively (Pan & Heitman, 1999, Robertson & Fink, 1998, Toda *et al.*, 1987). In *Candida albicans*, the catalytic isoforms Tpk1p and Tpk2p share positive roles in cell growth and hyphae formation, while they have distinct roles in hyphal morphogenesis, stress response and regulation of glycogen metabolism (Bockmühl *et al.*, 2001, Cloutier *et al.*, 2003, Sonneborn *et al.*, 2000). Meanwhile, numerous known filamentous fungi are found to possess only two distantly related PKA-C isoforms with varied functions (Banno *et al.*, 2005, Lee *et al.*, 2003, Ni *et al.*, 2005, Schumacher *et al.*, 2008). In most plant pathogenic fungi, deletion of one PKA-C isoform resulted in profound effects in virulence. For instance, of the two PKA catalytic subunits, only Adr1 kinase activity is essential for the dimorphic transition and pathogenesis in *Ustilago maydis* (Dürrenberger *et al.*, 1998). Similarly, only one PKA isoform plays a predominant role in other phytopathogens like *Colletotrichum trifolii*, *C. lagenarium*, *Botrytis cinerea* and *Setosphaeria turcica* that utilize appressoria to penetrate and infect the host; or in *Mycosphaerella graminicola* and *Verticillium dahlia*, that invade the host through stomata or other natural openings (Hao *et al.*, 2015, Schumacher *et al.*, 2008, Takano *et al.*, 2001, Yamauchi *et al.*, 2004, Yang & Dickman, 1999, Mehrabi & Kema, 2006, Tzima *et al.*, 2010).

The Rice Blast pathosystem has been extensively analysed at the molecular level, and serves as a model in the study of plant–fungus interactions (Liu *et al.*, 2013), and in tackling global food security (Nalley *et al.*, 2016). The cAMP/PKA signaling in *M. oryzae* plays an important role in surface sensing, appressorium morphogenesis, turgor generation, and in regulating plant infection (Li *et al.*, 2012, Yan & Talbot, 2016). Different components of the G-protein signaling such as the Gα, MagA, MagB or MagC, two Gβ (Mgb1 and Mgb2), a Gγ subunit, and the Rgs1 (regulator of G-protein signaling 1) have been characterized (Dean *et al.*, 2005, Fang & Dean, 2000, Liu *et al.*, 2007, Liu & Dean, 1997, Nishimura *et al.*, 2003, Ramanujam *et al.*, 2012). Anchoring and trafficking of G-protein signaling components on late endosomes endows *M. oryzae* with the ability to specifically activate, integrate and achieve modularity and spatio-temporal control of signaling responses critical for pathogenesis (Ramanujam *et al.*, 2013). Downstream of the G proteins, the adenylate cyclase Mac1 (that synthesizes cAMP), its suppressor Sum1, and the cAMP phosphodiesterases have been characterized too (Adachi & Hamer, 1998, Choi & Dean, 1997, Ramanujam & Naqvi, 2010, Zhang *et al.*, 2011). Mutants disrupted in the catalytic subunit gene *CPKA* exhibit normal growth and conidiation, but show delayed appressorium formation and loss of pathogenicity, which results from the defects in appressorial function (Mitchell & Dean, 1995, Xu *et al.*, 1997). Recently, we showed that loss of the regulatory subunit of PKA (*RPKA*) results in complete loss of pathogenicity; and a suppressor mutant that partially restores the pathogenicity in *rpk*AΔ represents a point mutation in the *CPKA* locus (Selvaraj *et al.*, 2017). These studies confirm a crucial role for cAMP/PKA signaling in the development and pathogenicity of *M. oryzae.* However, the second catalytic subunit of PKA, Cpk2, has been predicted but not characterised thus far.

In this study, we analysed the functions of *CPK*2 and further created a *cpk*A*cpk*2 double mutant to study the complete effects of PKA signaling on the pathogenicity of *M. oryzae*. We show that although the Cpk2 activity is largely redundant in function to that of CpkA, both catalytic subunits act in concert to regulate hyphal growth and play overlapping roles in conidiation and appressorium formation in *M. oryzae*. Importantly, these processes are dependent on Cpk2, since deletion of *CPK*2 removes even the residual virulence associated with loss of *CPK*A. The expression of *CPK*2 under the *CPK*A promoter, or the swapping in of *CPK*2 coding region for *CPK*A, restored the pathogenicity in *M. oryzae cpkA* null mutant. Furthermore, Cpk2-GFP localizes to a different compartment, the nucleus, and is independent of CpkA. Taken together, this study underscores the importance of cyclic AMP PKA signaling in the pathogenesis of *M. oryzae*.

## RESULTS

### Identification and gene-deletion analysis of *CPK2* in *M. oryzae*

The sequence of the *M. oryzae* genome (http://fungi.ensembl.org/ *Magnaporthe_oryzae*/Gene/Summary) revealed an open reading frame (ORF) that encodes a catalytic subunit of PKA, *CPK2* (contig 6–2325: coordinates 2566–1350), which was different from *CPKA*. *CPK2* encodes a member of the class II PKA subunits found uniquely in filamentous fungi (Schumacher *et al.*, 2008), the function of which is not yet known in *M. oryzae*. MGG_02832 (GI: 2682385) showed the presence of three exons spanning a 1474 bp ORF, and is predicted to encode a 408-amino acid polypeptide (XP_003720907.1; protein reference). The Cpk2 protein contains a typical serine/threonine kinase domain as well as a C-terminal AGC domain, which is representative of a large family of kinases and conserved in numerous PKA catalytic subunits (Pearce *et al.*, 2010). *M. oryzae* Cpk2 shows 40 to 53% amino acid identity to yeast PKAs (Toda *et al.*, 1987) and high amino acid identity with other class II PKAs of *A. fumigatus* (Liebmann *et al.*, 2004), *B. cinerea* (Schumacher *et al.*, 2008), *U. maydis* (Dürrenberger *et al.*, 1998) - for a detailed phylogenetic alignment refer (Schumacher *et al.*, 2008). The CpkA and Cpk2 proteins of *M. oryzae* share approximately 48% sequence identity, with the highest degree of divergence within the N-terminal region. The highly conserved protein kinase domain of Cpk2 extended from 75-350 amino acids and the nucleotide binding site LGTGFARV (81-89 aa) differed by 1 amino acid from the conserved motif of CpkA, whereas the active catalytic domain RDLKPEN (203-215 aa) was identical (Fig. 1).

**Fig. 1.**
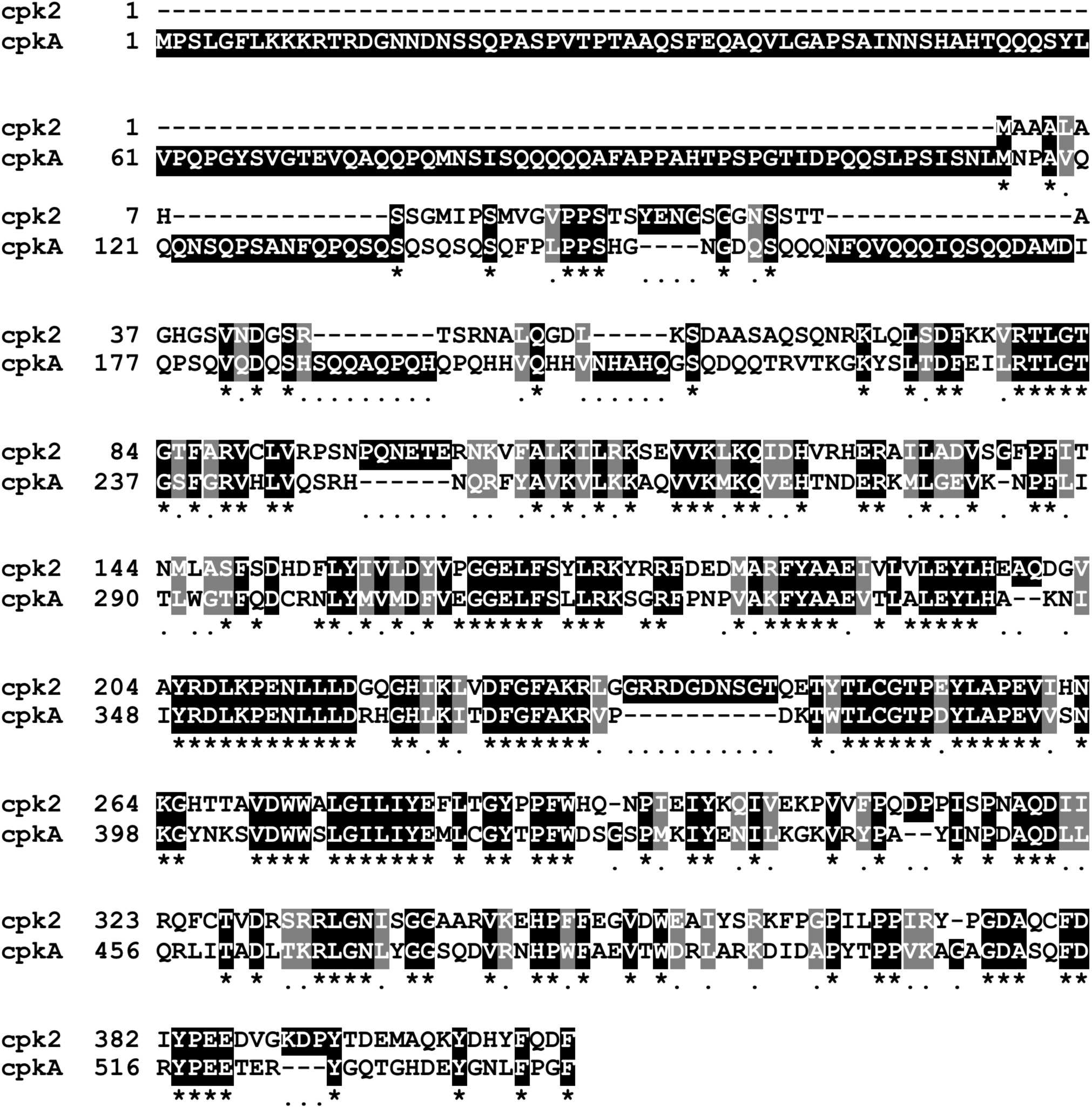
Amino acid sequence alignment for the PKA-C subunits, CpkA and Cpk2, from *M. oryzae*. The protein sequences of CpkA and Cpk2 showing identities (shaded black) and similarities (gray background). Alignment was carried out using ClustalW of the MegAlign software, a program of the Lasergene package (DNASTAR) using default parameters; and shaded with BOXSHADE 3.2.1 (www.ch.embnet.org/software/BOX_form.html).

We carried out gene-deletion analyses to determine the relative contributions of each catalytic subunit of PKA, *CPK*A and *CPK*2, to the pathogenicity of *M. oryzae.* In addition, a strain deficient for both *CPK*A and *CPK*2 was generated illustrating that cAMP-PKA signaling is not essential for viability in *M. oryzae*. The transformants were confirmed through Southern blotting and locus specific PCR (Fig. S1), and in all cases at least two independent strains were characterized in detail.

### Cpk2 and CpkA are required for vegetative growth and conidiation in *M. oryzae*

The colony morphology and the radial growth of the individual *cpk*AΔ and *cpk*2Δ strains were indistinguishable from the WT, while the *cpk*AΔ*cpk*2Δ showed reduced radial growth producing small colonies with fluffy aerial growth (Fig. 2A and C). Although *CPKA* is dispensable for vegetative growth and conidiation, it regulates appressorium formation and function, with the *cpk*AΔ strain displaying long germ tubes and delayed appressorium formation (Mitchell & Dean, 1995, Xu *et al.*, 1997). No apparent changes were observed in conidiation in *cpk*AΔ and *cpk*2Δ conidiation compared to the WT. Based on the *cpk*2Δ phenotypes, we inferred that Cpk2 functionality could be elucidated better in the context of the loss of CpkA activity. Accordingly, deletion of *cpk*2 in *cpk*AΔ (*cpk*AΔ*cpk*2Δ) produced very few conidia (almost 90% reduction) compared to *cpk*AΔ or *cpk*2Δ or the WT. Further analysis revealed that conidiation was significantly delayed in the *cpk*AΔ*cpk*2Δ. The WT or the individual PKA mutants produced conidia within 24 h of light exposure, while the *cpk*AΔ*cpk*2Δ mutant didn’t initiate conidiation even after 48 hpi. At 7 to 10 d of exposure to light, the double mutant produced about 10-fold lesser conidia than the wild type (Fig. 2B and C). However, the conidia produced were three celled and with no apparent abnormalities in shape or size, thus implying that complete loss of PKA activity would adversely affect asexual development. We conclude that cAMP-PKA activity is essential for conidiation in *M. oryzae;* and CpkA and Cpk2 possess overlapping roles in regulating such metabolic activation and initiation of asexual reproduction therein.

**Fig. 2.**
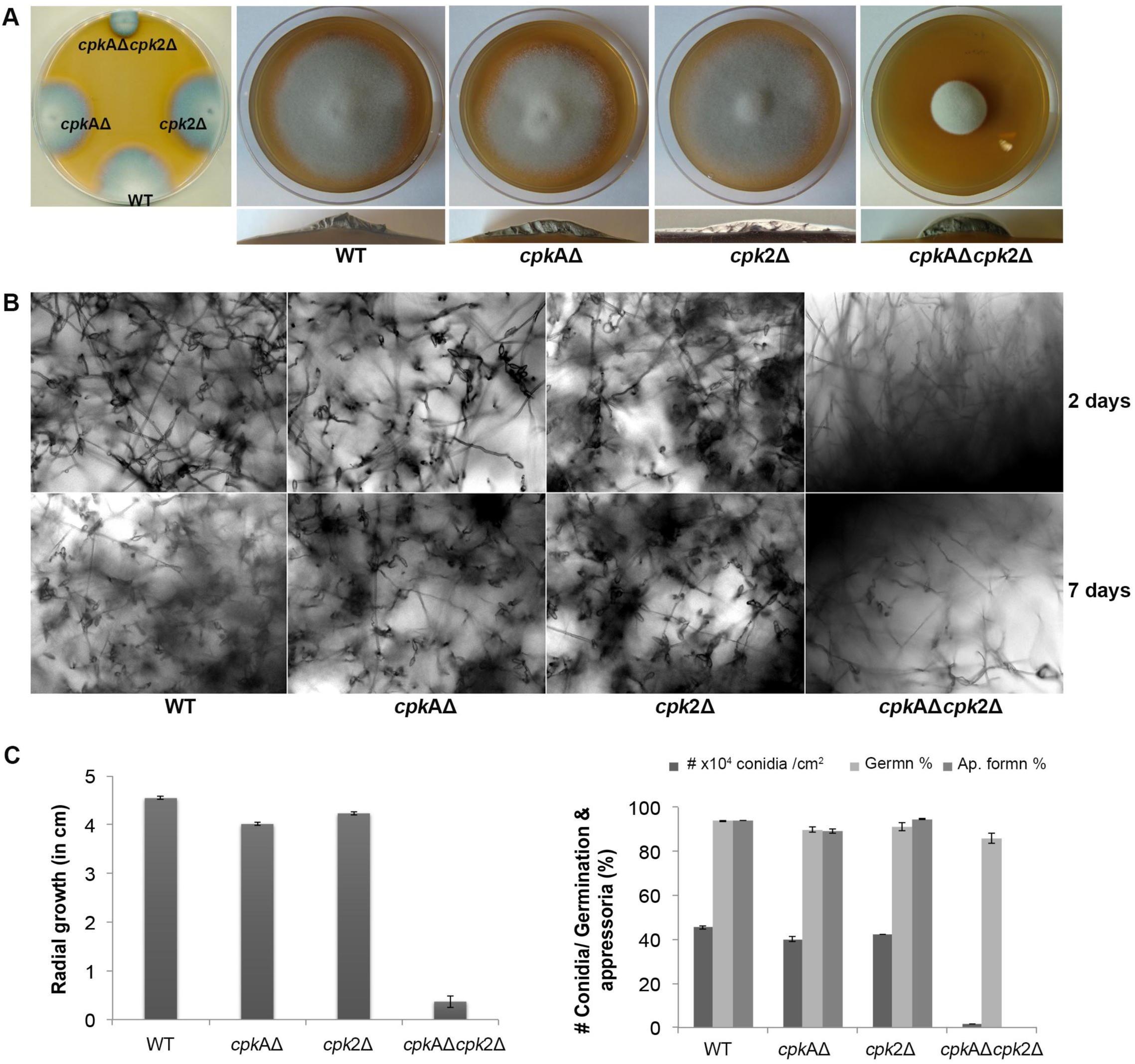
Cpk2-mediated cAMP-PKA signaling is necessary for proper vegetative and asexual development in *M. oryzae* (A) Radial and aerial hyphal growth of the wild type and the indicated PKA-C mutant strains. Mycelial plugs inoculated on PA medium was cultured in the dark at 28 °C for 5 days. The left panel shows the comparative radial growth of individual CPK mutants *cpkA*Δ, *cpk2*Δ and *cpkA*Δ*cpk2*Δ with the wild type B157. The radial and cross-sectional view for the aerial hyphal growth of WT and the mutants are shown (right). (B) Bright field micrographs showing the the conidiation at 48 h and 7 days post photoinduction. Individual *cpk*AΔ and *cpk*2Δ produced conidia normally as the WT at 48 h, while *cpk*AΔ*cpk*2Δ showed very few conidia even at 7 dpi. Scale bar = 10 μm. (C) Bar graphs showing the difference in radial growth (left) and the quantification of conidiation and appressorium formation in WT and PKA-C mutants. Values represent mean ± S.E of three independent replicates with approximately 200 conidia assessed per experiment.

### *CPK2* plays a significant role in appressorium formation in *M. oryzae*

The *cpk*AΔ showed a significant delay in appressoria formation on hydrophobic/inductive surfaces, and the appressoria produced were smaller with long germ tubes. The *cpk*2Δ produced normal appressoria indistinguishable from WT. Interestingly, the *cpk*AΔ*cpk*2Δ conidia germinated normally, but were incapable of appressoria formation. Germ tubes produced by *cpk*AΔ*cpk*2Δ conidia were very long, curled, showed periodic clockwise twists, and failed to elaborate appressoria even at 32 hpi (Fig. 3A and 3B). Furthermore, exogenous addition of cAMP did not restore appressorium formation in the *cpk*AΔ*cpk*2Δ mutant (Fig. 3C). Like the WT, the *cpk*AΔ and *cpk*2Δ elaborated appressoria on non-inductive surfaces in response to exogenously added cAMP. However, addition of cAMP to the conidia of *cpk*AΔ*cpk*2Δ on non-inductive surfaces did not induce such appressorium differentiation, revealing that *cpk*AΔ*cpk*2Δ is non-responsive to the cAMP stimulus (Figure 3D). Appressorium morphogenesis is tightly regulated by the cell cycle, with DNA replication and one round of mitosis being essential for the initiation of appressoria in *M. oryzae* (Li *et al.*, 2017, Saunders *et al.*, 2010, Veneault-Fourrey *et al.*, 2006). We inferred that the defect in appressorium formation in the *cpk*AΔ*cpk*2Δ is likely due to the inability to sense and/or respond to cAMP in addition to defects in surface sensing and adhesion. Based on the delay in appressoria formation in *cpkA*Δ, and the inability to initiate appressoria in *cpk*AΔ*cpk*2Δ, we conclude that *CPK2* plays an important role in surface sensing and appressorium formation in *M. oryzae*.

**Fig. 3.**
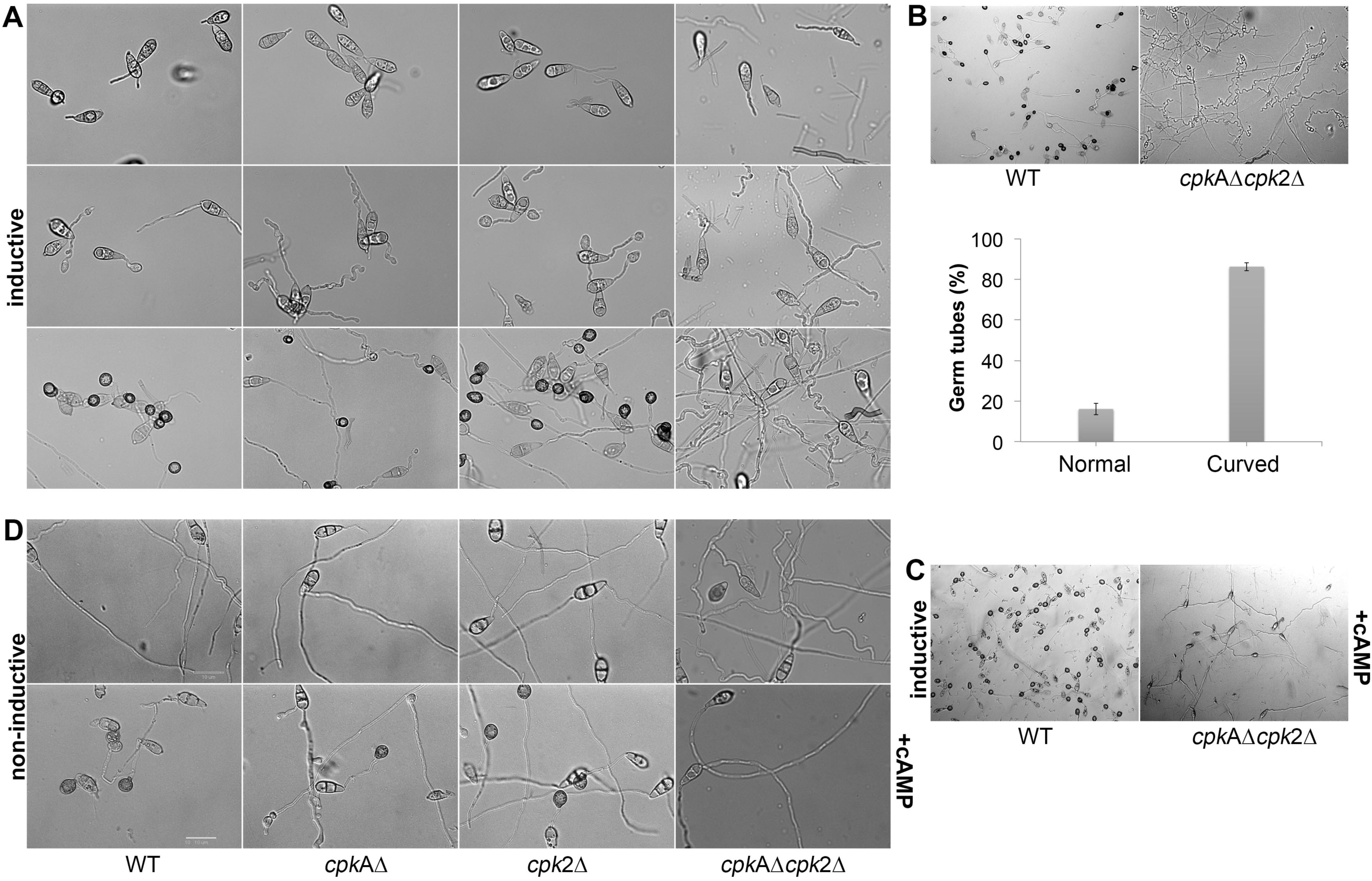
Cpk2 plays a crucial role in appressorium formation. Bright field micrographs of the appressoria formed by WT and CPK mutants on inductive - hydrophobic (A) and non-inductive - hydrophilic surfaces (D) at different time points. (A) The *cpk*AΔ*cpk*2Δ did not produce appressoria on inductive surface even at 24 hpi, while *cpk*AΔ was significantly delayed in appressorium formation. (B) The *cpkA*Δ*cpk2*Δ produced very long, wavy germ tubes without forming appressoria, and the germ tubes showed curled/zig-zag growth unlike the straight/linear growth mode in the WT. Different phenotypes of the germ tube formed by cpkAΔ*cpk*2Δ were quantified and represented as a bar graph. (C) Addition of exogenous cAMP (10 mM) to the conidia suppressed the wavy growth pattern, but failed to restore the appressoria formation in the *cpk*AΔ*cpk*2Δ. (D) Addition of exogenous cAMP (10 mM) to the conidia restored the appressorium formation in WT and *cpk*AΔ, *cpk*2Δ on non-inductive surface while *cpk*AΔ*cpk*2Δ was non-responsive to such exogenous cAMP Scale bar = 10 μm.

To further clarify the role of *CPK*2, we added the cAMP-PKA inhibitor KT5720 to the conidia on coverslips and checked appressorium formation at 24 h. Addition of KT5720 delayed appressorium formation in the WT, but had no effect on appressorium morphology. The appressorium formation on inductive surfaces was abolished, when the minimal PKA activity in *cpk2*Δ mutant was further blocked by the PKA inhibitor (Fig. S2), Thus strengthening the importance of *CPK2* in appressorium formation. However, the inhibitor concentration used in our experiment was not sufficient to completely block the PKA activity in the WT strain. We conclude that albeit redundant, the second catalytic cAMP-PKA subunit, Cpk2, plays an important role in appressorium formation in the rice blast fungus.

### *CPKA* and *CPK2* are involved in regulation of PKA signaling and intracellular cAMP levels

PKA activity was undetectable in the total protein extracts from the mycelia of *cpk*AΔ or *cpk*AΔ*cpk*2Δ, whereas the WT clearly showed the cAMP-dependent PKA activity (Fig. 4A). PKA activity could still be detected in the mycelial extracts of *cpk*2Δ but was very low. The *cpk*2Δ produced only about 50% PKA activity *in vitro* compared to the wild type *M. oryzae* (Fig. 4B) indicating that there is no compensatory increase in Cpk2 activity in the absence of *CPKA.* However, the function(s) of Cpk2 cannot be ascertained on the basis of its enzyme activity and the regulatory interactions between the two isoforms cannot be ruled out (Ni *et al.*, 2005). Hence, to determine the expression levels of *CPKA* and *CPK2* and to check if deletion of *CPKA* affects the expression of *CPK2* or vice versa, we extracted the RNA from WT, *cpkA*Δ and *cpk2*Δ and carried out qRT-PCR. In WT, albeit having a similar expression pattern, the level of *CPK*2 was comparatively lower than *CPKA* at all the time points (Fig. 4C), which indicates the likely reason for lower PKA activity in the *cpk2*Δ mutant. *CPKA* and *CPK*2 were expressed in mycelia and aerial hyphae at comparable levels, and hence the *cpk*AΔ*cpk*2Δ was highly impaired in radial and aerial growth. The expression of both isoforms increased in the appressoria at 8 h, which could be responsible for the indicated high levels of PKA activity during appressorium formation (Kang *et al.*, 1999). Based on the higher levels of transcription of the PKA isoforms in conidia and appressoria, we infer their functional importance in conidia and appressoria formation and pathogenicity. However, deletion of *CPKA* or *CPK2* had only a minor effect (less than two-fold) on the expression of the other PKA isoform at the time points analyzed (Fig. 4D). Hence, we rule out the possibility of co-transcriptional regulation between *CPKA* and *CPK2* in *M. oryzae*.

**Fig. 4.**
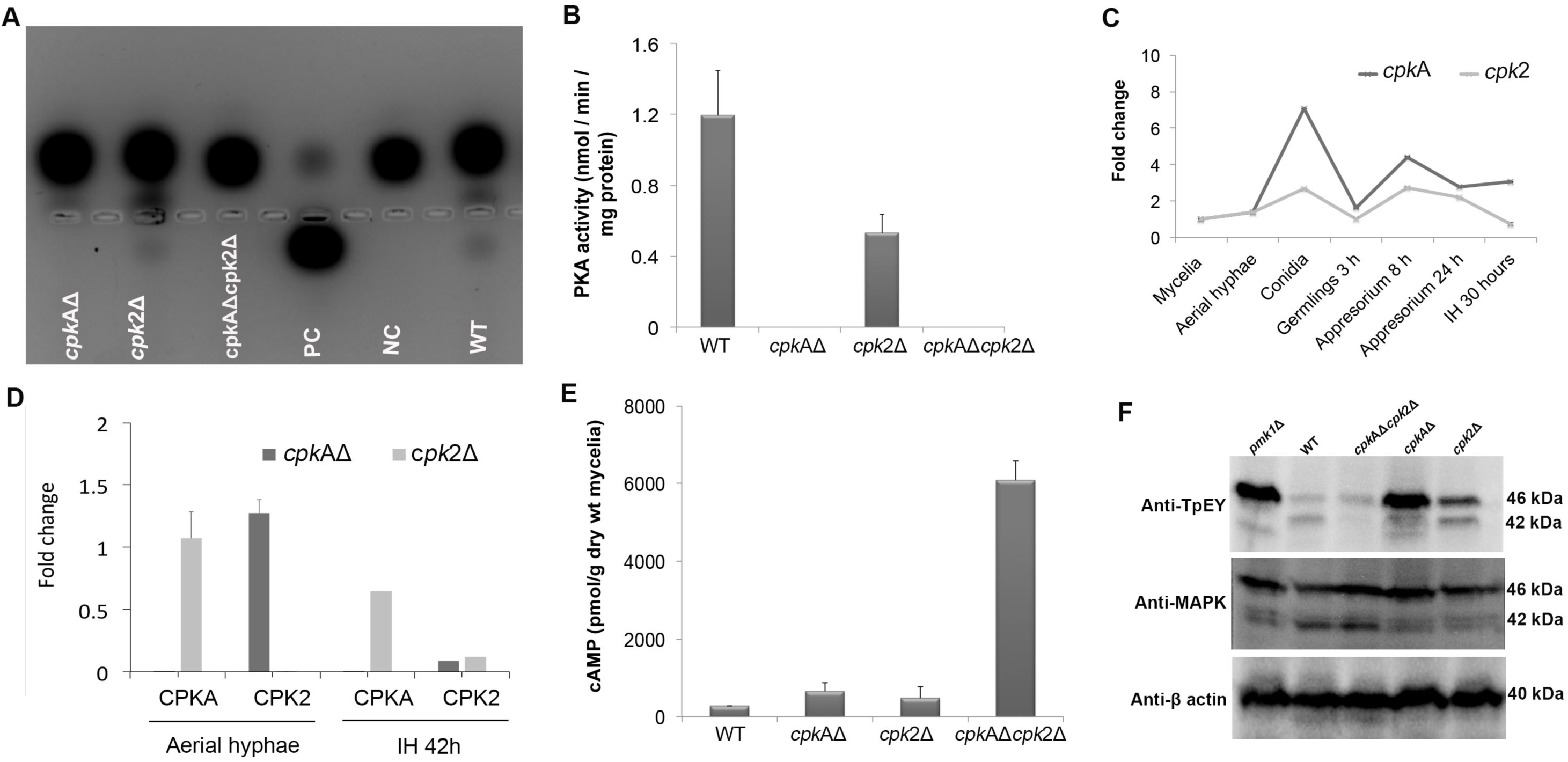
PKA activity regulates the intracellular cAMP levels, and the Pmk1-MAPK signaling in *M. oryzae*. (A) PKA enzyme activity was monitored by gel electrophoresis showing the migration of phosphorylated substrate towards the anode. PC-positive control; NC-negative control. PKA activity was analysed in the total protein extracts from the frozen mycelia of WT and the indicated CPK mutants. The results were consistent with repeated experiments. (B) Bar graph showing the PKA activity in WT and the listed PKA-C mutants. Each value represents a mean ± standard error of three replications. (C and D) Graphical representation of the fold change in the transcript/mRNA levels of *CPKA* and *CPK*2 in WT and the indicated mutants at the different stages of vegetative and pathogenic development. Reverse transcriptase RT-PCR was conducted on total RNA extracted from the mycelia and the infection structures mentioned. Fold change in gene expression was calculated from the average of three independent measurements, with β-Tubulin of *M. oryzae* as internal control and normalized to unit mycelial biomass. Error bars represent S.E. The experiment was repeated twice with three replicates each. (E) Quantification of the intracellular cAMP levels in the mycelia of WT and mutant strains. Values from two biological replicates with two replications for each individual sample were analysed and the mean value is presented. (F) Western blots showing the phosphorylation of Pmk1 (42 kDa) and Mps1 (46 kDa) MAPK in the WT and indicated PKA-C mutants. Total proteins were extracted from the mycelium of WT and mutants and analysed with the indicated antibodies. The *pmk1*Δ served as a negative control and the experiment was repeated twice.

The PKA-C mutants showed increased intracellular cAMP levels compared to the WT. The *cpk*AΔ had higher cAMP levels than *cpk*2Δ, consistent with the predominant role for CpkA in overall PKA activity/function. However, *cpk2* deletion either in WT or *cpk*AΔ led to an increase in cAMP concentration indicating that *cpk2* may also act as a cAMP effector though it is not essential in WT. The cAMP increased to very high levels in *cpk*AΔ*cpk*2Δ, indicating an additive effect of the loss of both catalytic subunits of cAMP-PKA (Fig. 4E). We infer that cAMP-PKA functions to limit the intrinsic cAMP to a threshold/moderate level to maintain normal cellular functions; or conversely, the activation of PKA dampens the intracellular cAMP. We conclude that Cpk2 acts in concert with CpkA to regulate the overall accumulation and dynamics of cAMP signaling in *M. oryzae*.

### Pmk1 MAPK phosphorylation is affected in the cAMP-PKA mutants

The non-responsiveness for cAMP in *cpk*AΔ*cpk*2Δ that resembles the *pmk*1Δ phenotype (Kou *et al.*, 2016, Xu & Hamer, 1996) led us to assess whether the Pmk1 MAPK activation is compromised in the PKA-C mutants. Therein, we assayed the phosphorylation of Pmk1 with the anti-TpEY specific antibody that detects the phosphorylation of both Pmk1 and Mps1 MAPKs (Zhao et al., 2005). In WT, a band of 42 and 46 kDa indicating the phosphorylation of Pmk1 and Mps1 were observed, while in *pmk1*Δ only the Mps1 phosphorylation was evident (Fig. 4F). Weak phosphorylation of Pmk1 was observed in *cpk*AΔ or *cpk*2Δ whereas such activation was completely absent in the *cpk*AΔ*cpk*2Δ. The level of Mps1 phosphorylation increased in *cpk*AΔ or *cpk*2Δ compared to the WT, but similar to *pmk1*Δ indicating that Mps1 is likely hyperactive in response to/or to compensate for the cell wall defects associated with loss of Pmk1 (Zhao *et al.*, 2005). These results showed that signaling through Pmk1 may be impaired but not completely blocked in *cpk*AΔ or *cpk*2Δ, but an additive effect is observed in the double mutant wherein the total loss of Pmk1 activation is likely responsible for the observed defects in appressorium formation.

### cAMP-PKA signaling and pathogenesis of *M. oryzae*

We tested the cAMP-PKA mutant strains for their ability to cause blast disease lesions in barley and rice. The *cpk*AΔ and *cpk*AΔ*cpk*2Δ failed to elicit any visible blast symptoms on barley leaves, whereas the WT or *cpk*2Δ inoculation resulted in typical blast lesions on barley leaves. Wounding of rice or barley leaves with the micropipette tip helped the *cpk*AΔ to produce WT-like blast lesions in such abraded tissues (Fig. 5A). While *cpk*AΔ was still able to elicit necrosis on rice roots comparable to the WT or *cpk*2Δ, the *cpk*AΔ*cpk*2Δ strain did not produce any visible disease symptoms or necrosis on rice roots (Fig. 5B). Analyses of the invasive growth in rice leaf sheath revealed that *cpk*2Δ was pathogenic was able to penetrate the host plants (28 hpi), and grow invasively into the neighbouring cells similar to the WT (42 to 72 hpi). Consistent with previous results, the appressoria produced by *cpk*AΔ were impaired in penetration and could be observed on the surface of the rice leaf sheath at 28 and 42 hpi. The *cpk*AΔ*cpk*2Δ failed to produce appressoria on leaf sheath even at 42 hpi and hence was deemed completely non-pathogenic (Fig. 5C). As observed with barley leaf assays, the spray inoculation of conidia from WT or *cpk*2Δ produced typical blast lesions, while *cpk*AΔ remained non-pathogenic (Fig. 5D). As *cpk*AΔ*cpk*2Δ produced very few conidia, we were unable to carry out spray inoculation assays for the double mutant strain.

**Fig. 5.**
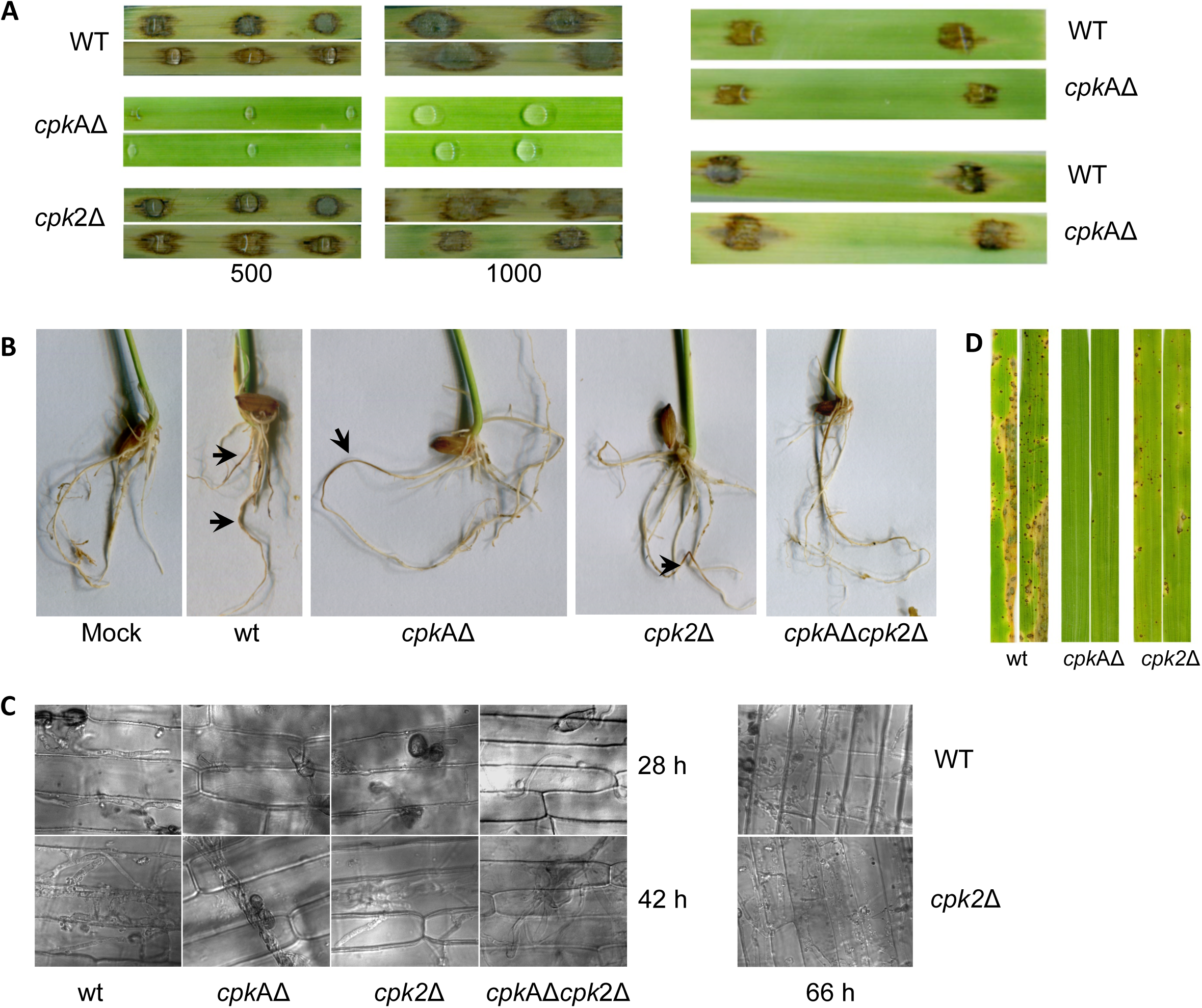
PKA signaling plays an essential role in the pathogenicity of *M. oryzae*. (A) Barley leaf infection assays with WT and the mutant strains. Conidia from WT or mutant strains were used to inoculate barley leaf explants and the blast disease symptoms/lesions were assessed 7 days post inoculation. Number of conidia used for each inoculation is indicated. For *cpkA*Δ, the abraded (wounded) leaves showed WT-like blast disease lesions (Right panel). (B) Rice root infection assays with WT and the PKA-C mutant strains. Surface sterilized rice seeds were allowed to germinate and grow on mycelial plugs of the wild type or the mutant strains; and necrosis/lesions were recorded after an incubation of 15 days. Mock indicates PA plugs without the fungal cultures. Arrows indicate necrosis/lesions on the roots. (C) Invasive growth of PKA-C mutants compared with WT when inoculated on rice leaf sheath. Bright field micrographs showing the invasive hyphal growth of WT and PKA-C mutants at the indicated time points. Scale bar = 10 μm. (D) Spray inoculation assays in rice to confirm the pathogenicity of WT and mutants. Conidia (1 X10^5^/ml, 5ml) from WT or PKA-C mutants were sprayed on four-week-old seedlings of rice cultivar CO39. Blast disease symptoms were assessed at 10 dpi.

### Swapping of the *CPKA* ORF with *CPK2*

In order to check if Cpk2 could functionally complement CpkA, we precisely replaced the *CPKA* coding sequence with the *CPK2* ORF, thus creating a genetic background that consequently lacks *CPKA*, but expresses *CPK2* under the *CPKA* promoter/regulon. The native *CPK2* remained unperturbed in such swapped strain. The resultant *CPKA*_Promoter_*CPK*2 strain showed WT-like vegetative growth, but displayed significantly reduced conidiation (Fig. 6A and B). The appressorium formation was delayed, and the resultant appressoria formed after prolonged germ tube growth were smaller in size similar to the *cpk*AΔ (Fig. 6C), again confirming that *CPKA* is required for proper appressorium formation. Furthermore, the PKA activity could not be detected in the total protein extracts from the mycelia of *CPKA*_Promoter_*CPK2* strain.

**Fig. 6.**
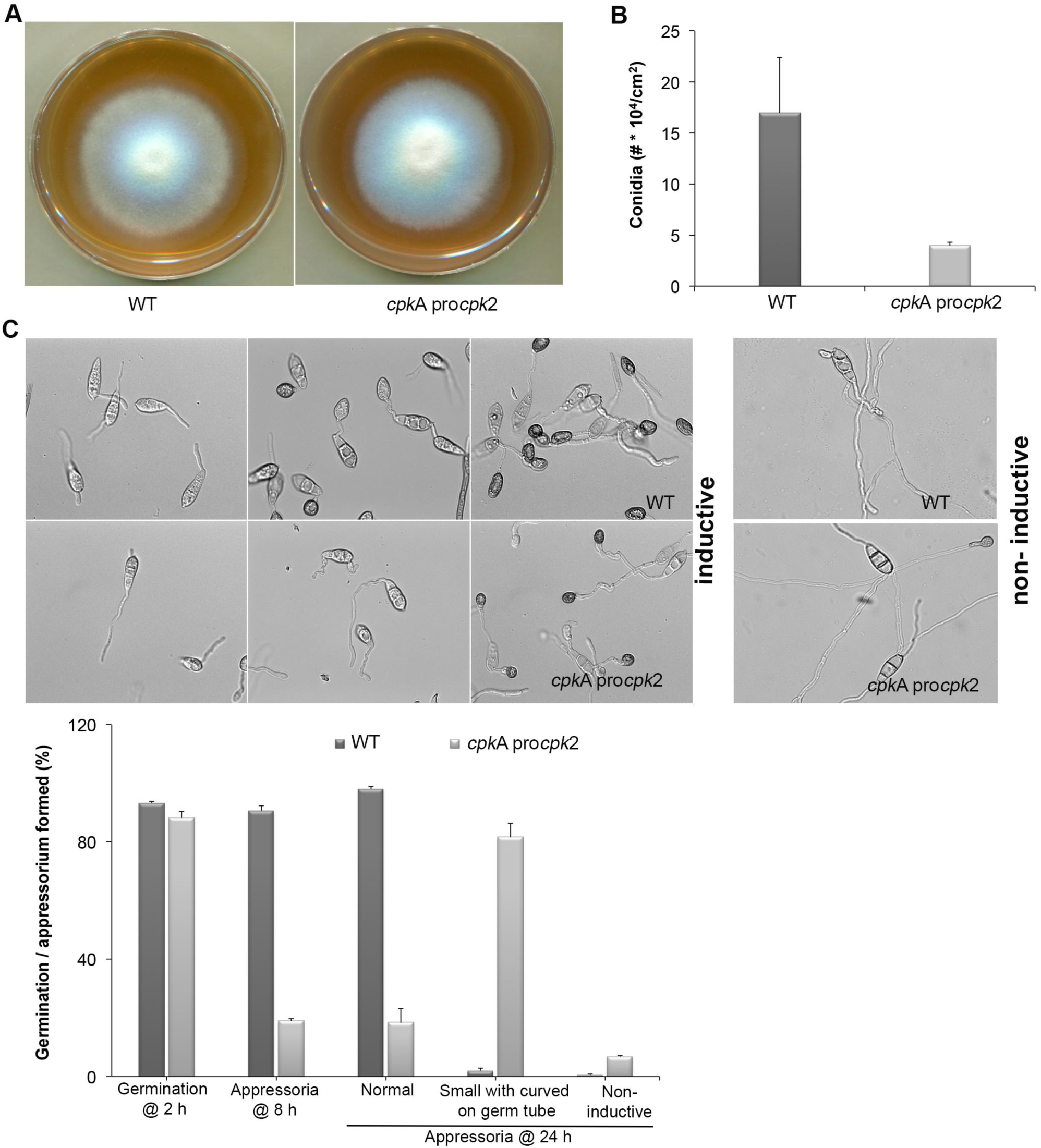
Cpk2 is able to partially compensate for the loss of CpkA (A) Radial growth of CPKApro*CPK2* compared with WT on prune-agar medium. The CPKAproCPK2 strain showed no defects in vegetative growth when compared to the wild type *M. oryzae*. (B) Quantification of conidiation revealed that substituting *CPK2* open reading frame for CPKA results in reduced conidiation implicating a major role of *CPKA* in conidiation. Each bar represents mean ± S.E of three independent replicates. (C) Appressorium formation on inductive (left) and non-inductive surfaces (lower right panel) and quantification of germination, appressorium formation by CPKApro*CPK2* compared with WT (upper right panel). Values represent mean ± S.E of three independent replicates using about 200 conidia per experiment.

Interestingly, the *CPKA*_Promoter_*CPK2* strain could produce blast lesions similar to the WT on rice leaves although the lesion size was smaller than the WT lesions on rice (Fig. 7A), thus indicating that the appressorial function is restored to some extent in the *CPKA*_promoter_*CPK2* strain unlike in *cpk*AΔ. The *CPKA*_Promoter_*CPK2* appressoria could penetrate the rice leaves, albeit delayed compared to the WT, and were able to successfully invade the host plants. However, the IH growth was compromised and the mutant strain remained restricted to the site of inoculation (Fig. 7B). Less than 20% of the *CPKA*_Promoter_*CPK2* appressoria could penetrate the rice plants at 28 h; and by 42 hpi more than 50% appressoria could penetrate and produce IH with only 10% capable of spread into the neighbouring cells. By 42 h, about 80% of the WT appressoria produced secondary IH (Fig. 7C). The *cpkA*Δ was able to produce a successful infection when inoculated through wounds as observed here and from previous reports. In contrast, the *CPKA*_Promoter_*CPK2* strain penetrated the rice sheath but was defective in IH growth. To check if the suppression of host penetration results from overexpression of *CPK2*, we analysed the transcript levels of *CPK2* in the *CPKA*_Promoter_*CPK2* strain compared to the WT. The results showed that *CPK2* is not highly expressed in this strain (Fig. S3) and hence the suppression of defects associated with loss of *CPKA* are likely due to the redundancy in Cpk2 function during pathogenic differentiation. To conclude, Cpk2 shares a potentially redundant function with CpkA, but is unable to compensate/complement the loss of *CPKA* activity in *M. oryzae*.

**Fig. 7.**
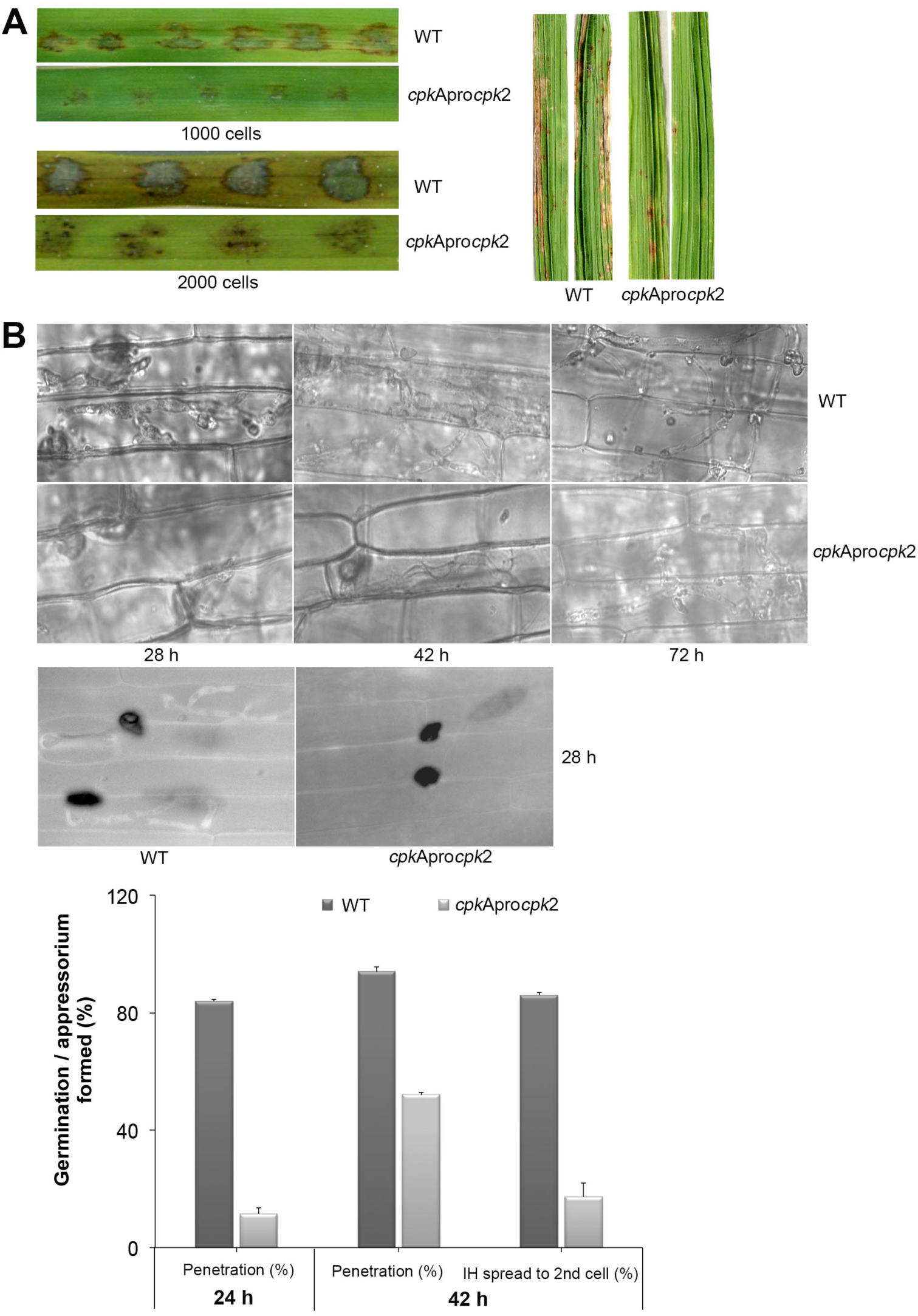
Substituting Cpk2 for CpkA partially suppresses the penetration and pathogenicity defects of *cpkA* deletion mutant. (A) Infection assays in rice with the conidia of CPKApromoter*CPK2* strain compared to the wild type. Conidia from WT and CPKApromoter*CPK2* were used to spot inoculate rice leaves, or sprayed on to seedlings of rice cultivar CO39 (right). The number of conidia inoculated on the detached barley leaves is indicated. Disease lesions were scored on 10 dpi for rice leaves and at 14 dpi for the spray inoculation assays. (B) Bright field micrographs showing the invasive hyphal growth of WT and CPKApromoter*CPK2* strain at the indicated time points (upper panel). Scale bar = 10 μm. The CPKApromoter*CPK2* showed impaired penetration and was defective in invasive hyphal growth. Micrographs showing host penetration by WT and CPKApromoter*CPK2* at 28h stained with aniline blue to show the callose deposition (lower left panel). Bar graph showing the host penetration ability and the percentage of appressoria that formed primary and secondary invasive hyphae from the WT and CPKApro*CPK2* at the indicated time points (lower right panel).

### Cpk2-GFP localizes to dynamic cytoplasmic vesicles, and the nucleus

To analyse the subcellular localization of Cpk2 during asexual and pathogenic phases of *M. oryzae*, the Cpk2 was fused with GFP at its C terminus. The expression from the native Cpk2 promoter was too weak to observe proper epifluorescence (Fig. S4). Hence, the GFP-Cpk2 fusion was expressed under the control of the constitutive Histone H3 promoter (*pH3GFP*-*CPK2*). The *in vivo* functionality of the fusion protein was verified through rigorous analysis of several phenotypes, and the aforementioned modified strains were found to be comparable to the parental untagged strain in all aspects of growth and pathogenicity. A double-tagged strain, *CPK2*-*GFP CPKA*-*mCherry*, was generated to examine the colocalization (if any).

The Cpk2-GFP was highly expressed in the vegetative stage i.e. in mycelia/ aerial hyphae on PA medium, compared to conidia and related structures therein (Fig. S4 -A, B). Staining with the Hoechst dye confirmed the nuclear localization of Cpk2-GFP during the mycelia growth phase (Fig. S4-C). The constitutively expressed GFP-Cpk2 (*H3Pro*G*FP*-*CPK2* strain) showed a similar localization pattern i.e. remained nuclear and cytoplasmic during mycelial growth. Nuclear localization of GFP-Cpk2 was clearly evident in conidia, and as cytoplasmic vesicles in the terminal cell of the conidia and also in germ tubes (Fig. 8, also see Fig. S5). The GFP-Cpk2 vesicles moved to the emerging appressorium at the hooking stage. In mature appressoria (24 h), GFP-Cpk2 showed a peri-nuclear vesicular localization, although the nuclear localization *per se* was not as prominent as in conidia (Fig. 8). In order to confirm the nuclear localization of GFP-Cpk2, the conidia from *H3ProGFP*-*CPK2* strain were co-stained with DAPI. GFP-Cpk2 colocalized with the DAPI signal, thus confirming the nuclear localization of Cpk2 therein (Fig. 9). The Cpk2-GFP vesicles co-localized with the CpkA-mCherry vesicles in the cytoplasm at different stages analysed. However, nuclear localization of Cpk2-GFP was not clear in this strain likely because of the weak Cpk2-GFP signal due to native expression, and/or due to masking by the stronger CpkA-mC expression (Fig. 10). We conclude that Cpk2 is compartmentalized in the nucleus, and its colocalization with CpkA is exclusive to the cytoplasmic vesicles. We infer that such intracellular localization facilitates RpkA interaction with both Cpk2 and CpkA, thus enabling robust cAMP-PKA enzyme activity/function to be regulated effectively in a compartmentalised manner. Lastly, the localization pattern clearly supports some special targets (and/or functions) for Cpk2 in activating the downstream cyclic AMP signalling in the nucleus during pathogenic differentiation in the rice blast fungus.

**Fig. 8.**
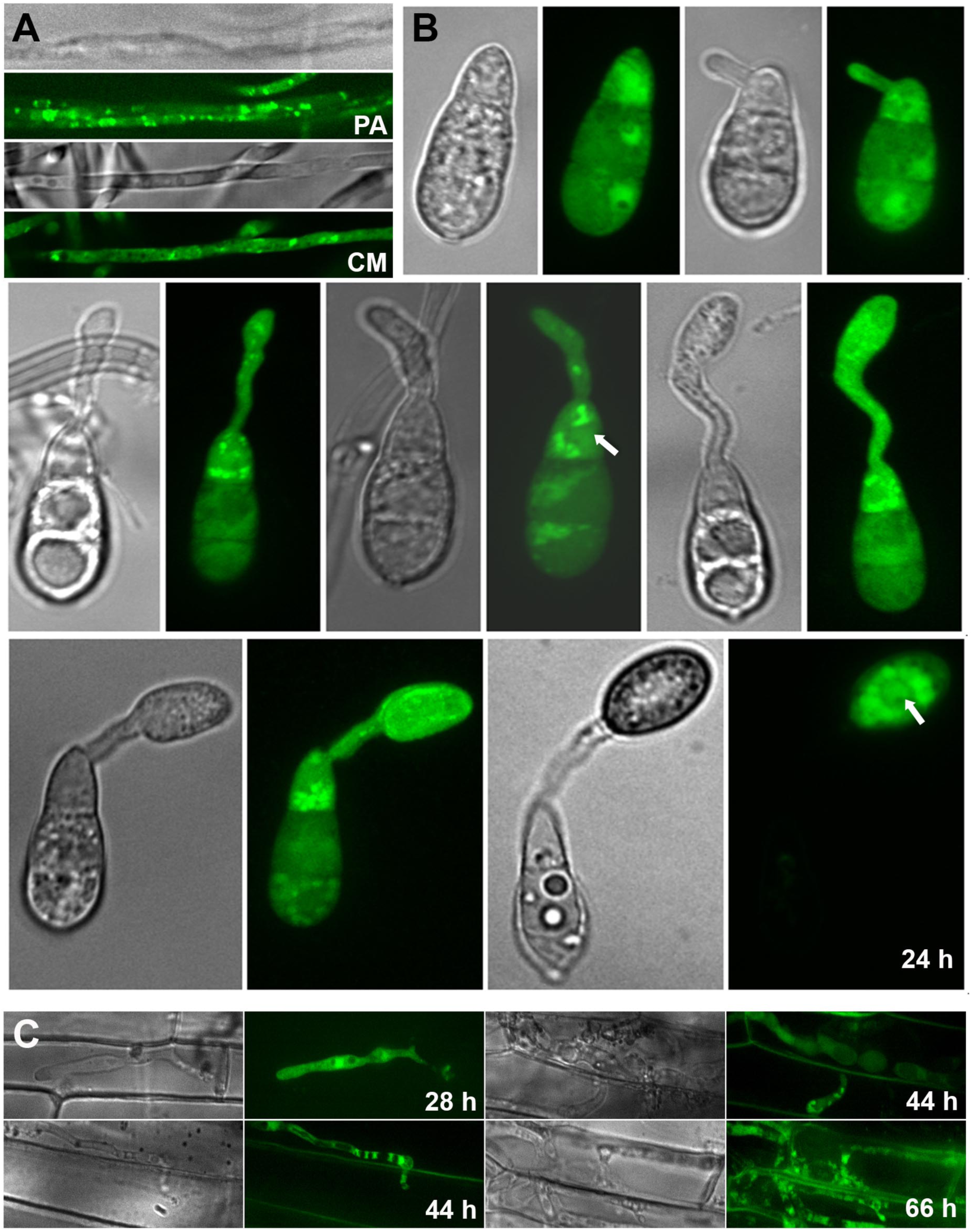
Subcellular localization of the GFP-Cpk2 at different stages of vegetative and asexual development in *M. oryzae*. (A) Vegetative hyphae and aerial structures from pH3GFP-CPK2-3 were imaged after growth on PA medium for 3 days. (B) Conidia inoculated on the inductive surface were subjected to time lapse analysis using a spinning disk confocal microscope. Bright field and the epifluorescent images were captured at the indicated time points using the requisite filters. Images are maximum intensity Z-projections of five confocal stacks, measuring 0.5 μm each. Scale bar is 10 μm. Arrows indicate the nuclear localization of GFP-Cpk2. (C) Subcellular localization of GFP-Cpk2 in the invasive hyphae formed in rice leaf sheath at 30 h after inoculation of CPKApro*CPK2*. Scale bar is 10 μm.

**Fig. 9.**
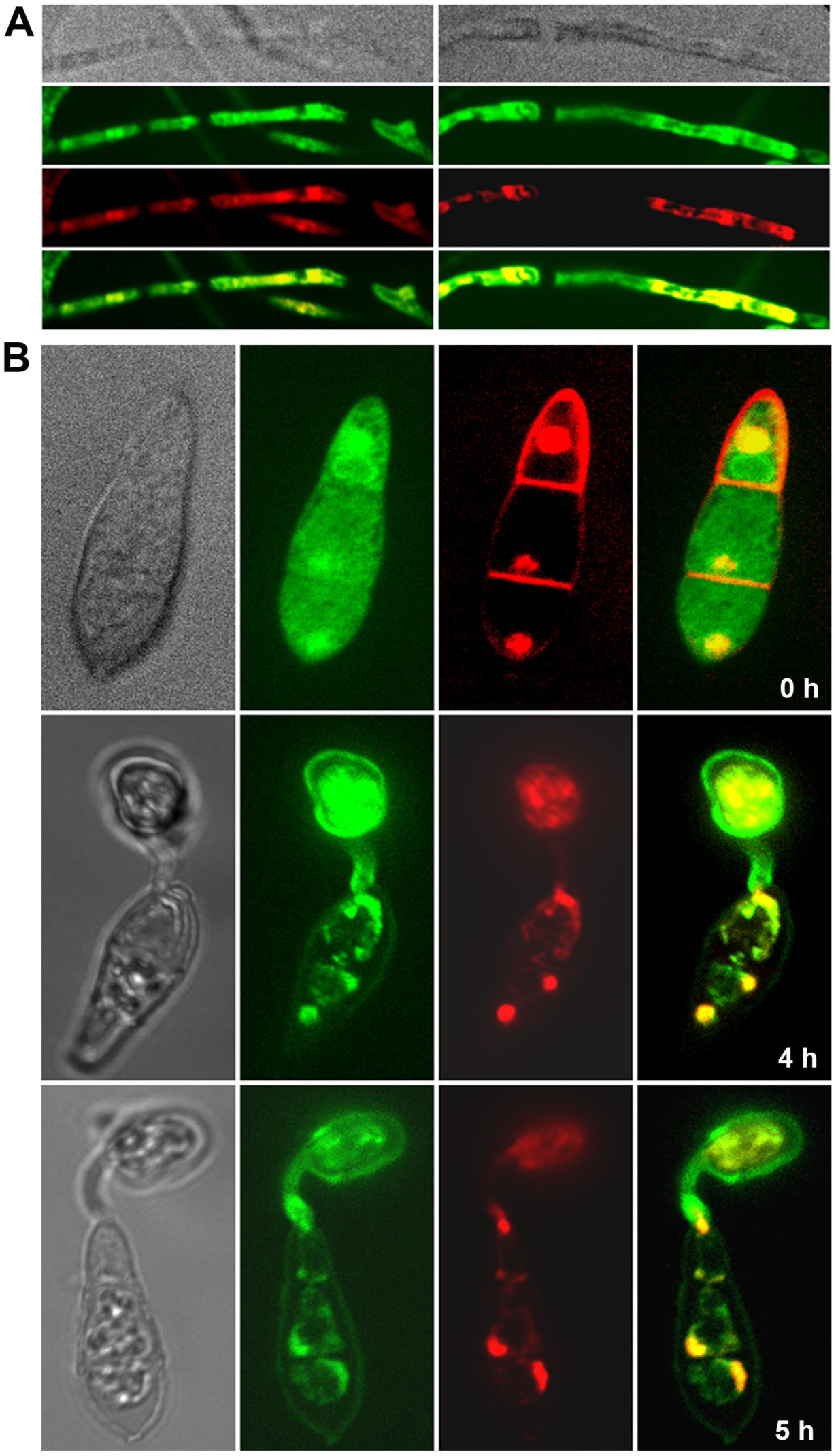
Confirmation of the nuclear localization of GFP-Cpk2. GFP-Cpk2 co-localizes with the nuclear stain DAPI at different structures as observed. Mycelia (A) or the conidia (B) were inoculated on coverslips and DAPI staining was carried out on fixed samples as described in methods. The co-localization (remains yellow in the merged panel) of GFP-Cpk2 (green) with nuclei/DAPI stained (pseudocolored red) is readily visible at the time points indicated. Images are maximum intensity Z-projections of five confocal stacks, 0.5 μm each. Scale bar equals 10 μm.

**Fig. 10.**
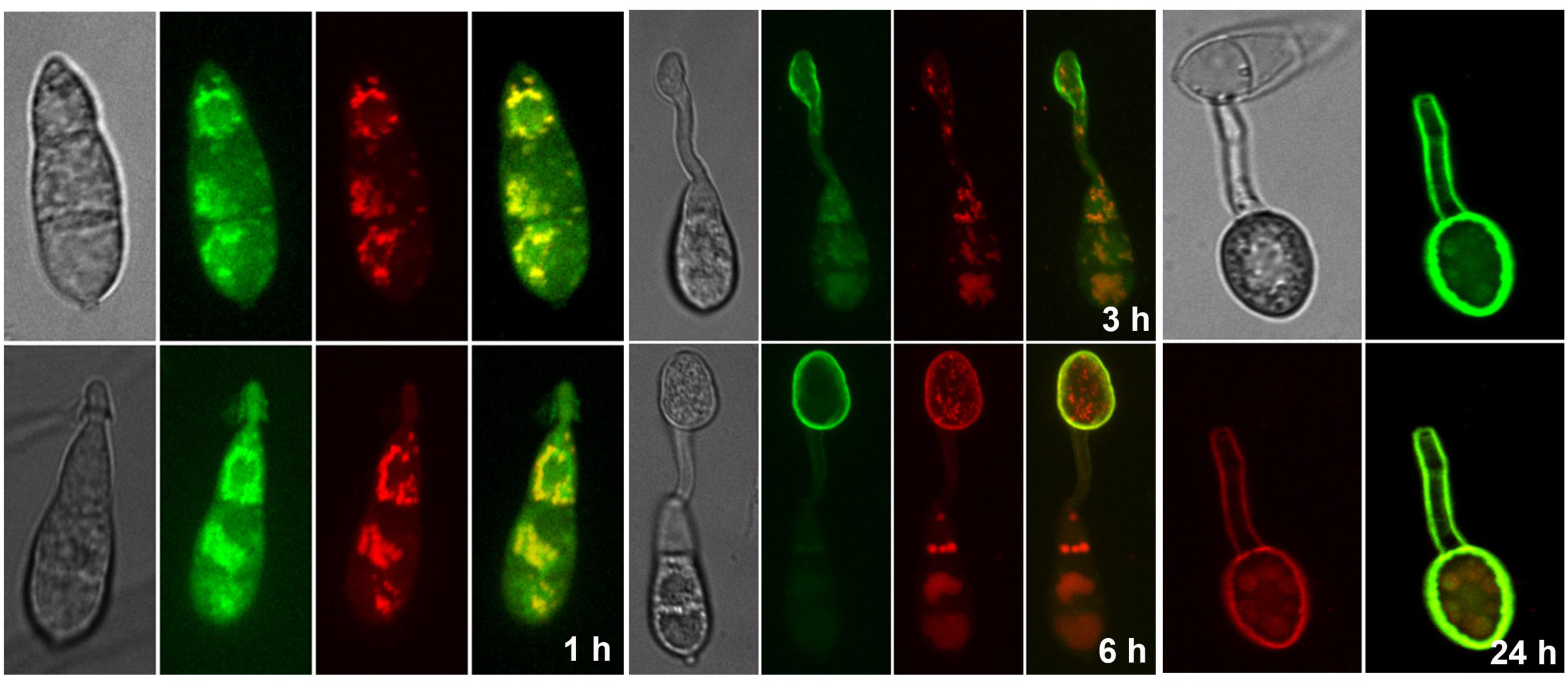
Co-localization of Cpk2-GFP with CPKA-mcherry. Cpk2-GFP cytoplasmic vesicles colocalize with CpkA-mCherry in the conidia and germ tubes. In the appressoria (24 h) a weak co-localization is evident in the vesicular structures within the perinuclear region.

## Discussion

A major challenge in deciphering the overall cAMP-PKA signaling is to fully understand the contribution(s) of both the catalytic subunits, CpkA and Cpk2, in *M. oryzae* pathogenicity. Most filamentous fungi examined to date, contain multiple isoforms of the catalytic cAMP-PKA subunit, with one such isoform playing a predominant role in growth and development. *M. oryzae* contains a divergent PKA-C isoform, Cpk2, but its contribution to growth and pathogenesis remained unexplored largely due to the indication that the *cpkAcpk2* double mutant is likely inviable (Choi & Xu, 2010). Despite its importance in regulating biological features and pathogenicity, the PKA catalytic subunits are not essential for cell viability in a number of fungal species (D’Souza & Heitman, 2001, Dürrenberger *et al.*, 1998, Fuller *et al.*, 2011). However, at least one TPK gene is required for cell viability and the *tpk1 tpk2 tpk3* triple mutant is not viable in *S. cerevisiae* (Toda et al., 1987). In *A. nidulans*, deletion of both *pkaB* and *pkaA* is lethal, though overexpression of *pkaB* can suppress some defects caused by Δ*pkaA*, indicating the overlapping roles of PkaA and PkaB (Ni *et al.*, 2005). Here, we showed that the cAMP-PKA signalling is not required for viability *per se* in *M. oryzae*, but is necessary for proper growth, conidiation and pathogenicity. This further adds to the previous findings that *MAC1* or *RPKA* are dispensable for viability but essential for pathogenic differentiation (Choi & Dean, 1997, Selvaraj *et al.*, 2017).

In *M. oryzae*, CpkA plays a role in appressorium morphogenesis and plant infection though it is dispensable for vegetative growth and conidiation, and is predicted to contribute the major PKA catalytic function (Mitchell & Dean, 1995, Xu *et al.*, 1997). Consistent with this observation, deletion of *CPK2* had no detectable effect on conidiation, appressorium formation or pathogenicity, thus making it redundant in function with *CPKA*. The enzyme activities and relative expression of the two isoforms in WT and mutant backgrounds revealed that cells do not compensate for the loss of one C isoform by overexpression of the other as in *S. cerevisiae* (MAZÓN *et al.*, 1993, Robertson *et al.*, 2000) and their interactions are certainly not regulated at the transcriptional level. However, we have shown here the functional capacity of Cpk2 to act in concert with CpkA in the regulation of vegetative growth and conidiation in *M. oryzae.* Appressorium induction upon proper surface sensing is a crucial step for pathogenesis in *M. oryzae*, and our results reflect the importance of Cpk2 function in this process. Our findings suggest that Cpk2, in addition to acting as an inducer of appressorium formation in concert with CpkA, also plays a regulatory role in suppressing appressoria biogenesis under unfavourable conditions such as on hydrophilic surfaces.

The defects of PKA-C null mutant (*cpkA*Δ*cpk2*Δ) resembled the phenotype of *mac1*Δ (Choi & Dean, 1997), despite the significantly higher accumulation of cAMP in this strain and also in individual mutants of the PKA-C subunits. In *S. cerevisiae* and *Cryptococcus neoformans*, it has been shown that the PKA-C regulate cAMP levels through a negative feedback loop by activating PDEs (Ma *et al.*, 1999). Preliminary results on cAMP levels in PKA mutants showed that such feedback inhibition on intracellular cAMP levels via PdeH occurs in *M. oryzae* too. Further, the downregulation of Pmk1 phosphorylation in the PKA-C mutants implies that the crosstalk between the cAMP and MAPK signaling might occur at the level of PKA-C. MoMsb2 and MoSho1proteins that function upstream from the *Pmk*1 cascade and have overlapping functions in recognizing various surface signals for *Pmk*1 activation and appressorium formation (Liu *et al.*, 2011). Although not characterized fully in this study, the defects in *cpkA*Δ*cpk2*Δ related to surface sensing and response, appear similar to the suppressor mutant phenotypes in the *CHM1*-deletion mutant (Li *et al.*, 2004). The molecular identity of such suppressor(s) of *chm1*Δ has not been ascertained yet. The *chm*1Δ also showed additional defects in hyphal growth, conidiation and appressorium formation, which could not be suppressed by exogenous cAMP (Li *et al.*, 2004).

Interestingly, we did not detect phosphorylation of the synthetic PKA substrate, kemptide, in any of the strains in which *CPKA* was deleted. Thus, Cpk2 is unable to serve as the primary PKA-C in *M. oryzae*. The Cpk2 overlap provides a basal level of PKA-C function to allow efficient vegetative growth and to maintain turgor for penetration of appressoria, but not inducible PKA function sufficient for conidiation and appressorium morphogenesis or proper IH growth. Nevertheless, the localization of Cpk2-GFP clearly implies some special functions for *CPK2* in *M. oryzae*. It is well recognized that compartmentalization of cAMP signaling allows spatially distinct pools of PKA to be differentially activated. PKA isoforms are anchored at specific intracellular sites by A-kinase anchoring proteins (AKAPs) in mammalian cells (Smith & Scott, 2006). However, AKAPs are not present in fungi. We showed earlier that cAMP signaling is compartmentalized in the nucleus and cytoplasm in *M. oryzae* (Ramanujam & Naqvi, 2010). The RpkA localizes to the nucleus whereas CpkA is present predominantly on cytoplasmic vesicles with the PKA holoenzyme being cytosolic (Selvaraj *et al.*, 2017). We infer that the nuclear pool of Cpk2-GFP is a consequence of its association with RpkA therein, and that this interaction drives the nuclear function of cAMP signalling in *M. oryzae*. The primary locale for CpkA being vesicular structures in the cytoplasm; and the RpkA and Cpk2 being in the cytoplasm and nucleus allows discrete cAMP-PKA modules that respond to discrete intracellular cAMP pools and subsequently modify specific target proteins. Furthermore, the compartmentalization of cAMP PDEs, the PdeH and PdeL to the cytoplasm and nucleus respectively (Ramanujam & Naqvi, 2010) also shows the importance of tailoring individual cAMP responses to precisely modulate the downstream signalling cascade.

In conclusion, proper PKA-C signaling is essential for the IH growth and pathogenicity and balanced activities of the CpkA and Cpk2 isoforms likely plays important roles in robust regulation of the infection process in *M. oryzae*. CpkA being the primary PKA, Cpk2 maintains important function(s) in regulating vegetative growth, conidiation and appressorium formation and also contributes to the spatial and temporal regulation of cAMP-PKA signaling in *M. oryzae*. CpkA and Cpk2 act in a redundant as well as parallel/specific manner to activate the downstream effectors of the cyclic AMP signalling and also Pmk1 MAPK during initiation and spread of the blast disease in rice. Future studies will focus on analysing the differential regulation and downstream targets of the two PKA-C isoforms in the rice blast pathosystem.

## EXPERIMENTAL PROCEDURES

### Strains, growth conditions and transformation

The *Magnaporthe oryzae* strain B157 obtained from the Directorate of Rice Research (Hyderabad, India) and its transformants/derived strains were routinely cultured on prune agar medium (PA) at 28°C for 7 to 10 days. Prune agar medium, Basal medium (BM) and complete medium (CM) were prepared as described previously (Selvaraj *et al.*, 2017). Assessment of growth, conidiation and appressorium formation were carried out as routinely. The cyclic AMP analog, 8-Br-cAMP (BioLog, Germany) was used at a final concentration of 10 mM with the stock (100mM) constituted in sterile water. *Agrobacterium tumefaciens*-mediated transformation (AtMT) was used to generate the transformants and CM with 250 mg/ml hygromycin (A.G. Scientific Inc, USA) or BM containing 40 mg/ml ammonium glufosinate or chlorimuron ethyl (sulfonyl urea, Cluzeau Info Labo, France) was used to select the fungal transformants. *Escherichia coli* strain XL1 was used for routine bacterial transformations, maintenance of various plasmids and *A. tumefaciens* AGL1 was used for T-DNA insertional transformations in this study. Requisite transformants were screened by Southern blot analysis and/or locus-specific PCR and in each case, two confirmed strains were selected for further observations.

### Nucleic acid manipulation and sequence analysis

The *CPK*2 orthologs were identified by searching the Genbank and fungal genome databases using the BLAST program (Altschul *et al.*, 1997) and multiple sequence alignments were carried out with ClustalW (Thompson *et al.*, 1994) and Boxshade (http://www.ch.embnet.org/software/BOX_form.html). Plasmid DNA extractions and genomic DNA extraction from the CM grown mycelium were carried out using standard kits; Geneaid High Speed Plasmid Mini kit and Yeast DNA purification kit (Epicenter Biotechnologies, USA) according to the protocols mentioned therein. The PCR primers used in this study are mentioned in Table S1. Nucleotide sequencing was performed using the ABI Prism big dye terminator method (PE Applied Biosystems). Southern blot analysis was performed by using the Enhanced chemiluminescent labeling and detection kit (Amersham Biosciences, RPN2108). Standard procedures were adopted for DNA restriction, agarose gel transformation and hybridizations for Southern blot (Sambrook *et al.*, 1989).

### Generation of *cpk*A and *cpk*2 deletion mutants, overexpression strains and GFP fusion constructs

To generate deletion mutants of *cpk*A and *cpk*2, gene replacement vectors encoding glufosinate ammonium resistance in pFGL97 or the hygromycin resistance in pFGL44 flanking the respective ORF were constructed using ligation PCR approach and then transformed to WT using AtMT. To get *cpk*AΔ*cpk*2Δ, the *cpk*2Δ construct was introduced into *cpk*AΔ strain. The *CPK2*-GFP in pFGL820 (encoding sulfonyl urea resistance gene cassette) was constructed by sequential cloning of the eGFP ORF, the last 1kb and the downstream fragment of CPK2 ORF to yield the final construct pCPK2-GFP-Trpc construct (Selvaraj *et al.*, 2017). The GFP-CPK2 overexpression construct was created by fusing the Moh3 promoter with the cpk2 ORF, sequentially cloned in to pFGL1010G which encodes sulfonyl urea resistance with an ilV locus which facilitate ectopic single copy integration. To construct a cpk2 ORF overlapping cpkA vector, the ORF of cpk2 was fused with the promoter of cpkA and ligated to pFGL880 (encoding sulfonyl urea resistance gene cassette) which already contained the *CPKA* 3’UTR.

### Protein isolation and western blot analysis

Total proteins (approximately 30 μg) from mycelia collected from 2 days old CM cultures extracted were separated on a 10 % SDSPAGE gel and transferred to nitrocellulose membranes for western blot analysis as described (Bruno et al., 2004; Liu et al., 2011). TEY phosphorylation of MAPKs was detected with the PhophoPlus p44/42 MAPK antibody kit (Cell Signaling Technology, Beverly, MA) according to the manufacturer’s instructions. A monoclonal anti-actin antibody (Sigma-Aldrich) was used to detect actin.

### Plant cultivar, growth and blast infection assays

Rice cultivar CO39 and barley cultivar Express susceptible to *M.oryzae* strain B157 were used for blast infection assays. Rice was grown at 80% humidity at 28 °C and Barley was grown at 60% humidity at 24°C (day) and 22 °C (night) with 12 h:12 h day:night cycles in a growth chamber. For plant infection assays, freshly harvested conidia at a concentration of 1 x 106 /ml in 0.2 % gelatin were used. Plant inoculation, incubation and lesion examination were conducted as mentioned previously (Ramanujam & Naqvi, 2010, Selvaraj *et al.*, 2017). Rice leaf sheath infection assay was performed as described (Kankanala *et al.*, 2007). Surface-sterilized rice seeds germinated and grown in direct contact with the fungal mycelial plugs were examined for black or browning lesions in the roots after two weeks to assess the root infection (Dufresne & Osbourn, 2001).

### Real Time qRT-PCR analysis

Total RNA was isolated from mycelia grown in CM for 2 days, conidia, germtubes or appressoria harvested from coverslips at different intervals and from rice leaves (CO39) infected with WT at 48 hpi using RNeasy Plant Mini kit (QIAGEN, USA) further treated by RNase-free DNase (Roche Diagnostics, Germany). The first strand cDNA was synthesised using the RevertAid first strand cDNA synthesis kit (K1622, Thermo Scientific) and used as template for qRT-PCR. qRT-PCR was performed on ABI 7900HT (Applied Biosystems, Foster City, CA, USA) using Power SYBR Green PCR Master Mix (Applied Biosystems, ThermoFisher Scientific) and the requisite primer sets for open reading frames for *cpk*A, *cpk*2 and β-tubulin (*TUB2*) (Table S1). All qRT-PCR reactions were conducted twice with three replications for each sample. The abundance of the gene transcripts was calculated by the 2^-ΔΔ^CTmethod with β-tubulin as the internal control.

### Assays for cAMP-dependent protein kinase A (PKA) and quantification of intracellular cAMP

PKA assay was performed using a nonradioactive cAMP-dependent protein kinase assay system fluorescent using the PKA model substrate, Kemptide (Promega, Madison, WI), following the manufacturer’s instructions and the samples were prepared as mentioned previously (Kang *et al.*, 1999, Selvaraj *et al.*, 2017). Protein concentrations in cell-free extracts were determined by the protein assay kit (Bio-Rad) according to the supplier’s instructions, and using BSA as a standard.

The samples for cAMP estimation were prepared as mentioned previously (Liu *et al.*, 2007, Ramanujam & Naqvi, 2010) and the assays was carried out using the cAMP Biotrak Immuno-assay System (Amersham Biosciences, NJ, USA) according to the manufacturer’s protocol.

### Microscopy, image analysis and processing

Staining with DAPI (diamidino-2-phenylindole; Sigma Aldrich, USA) was carried out essentially as described already (Patkar *et al.*, 2010, Ramanujam & Naqvi, 2010). Bright field and epifluorescence microscopy was performed with an Olympus IX71 or BX51 microscope (Olympus, Tokyo, Japan) using a Plan APO 100X/1.45 or UPlan FLN 60X/1.25 objective and appropriate filter sets. Images were captured with Photometrics CoolSNAP HQ camera (Tucson, AZ, USA) and processed using MetaVue (Universal Imaging, PA, USA), and Adobe Photoshop 7.0.1 (Mountain View, CA, USA). Time-lapse or live cell fluorescence microscopy was performed using a Zeiss Axiovert 200 M microscope (Plan Apochromat 1006, 1.4NA objective) equipped with an UltraView RS-3 spinning disk confocal system (PerkinElmer Inc., USA) using a CSU21 confocal optical scanner, 12-bit digital cooled Hamamatsu Orca-ER camera (OPELCO, Sterling, VA, USA) and a 491 nm 100 mW and a 561 nm 50 mW laser illumination under the control of MetaMorph Premier Software, (Universal Imaging, USA). Typically, z-stacks consisted of 0.5 μm-spaced planes for every time point. The maximum projection was obtained using the Metamorph built-in module. GFP excitation were performed at 491 nm (Em. 525/40 nm). Image processing and preparation was performed using Fiji (http://fiji.sc/wiki/index.php/Fiji), and Adobe Photoshop.

## Acknowledgements

We are grateful to the Fungal Patho-biology Group at TLL for discussions and suggestions. We thank M. Madhaiyan for help in organizing the figure panels using Photoshop. We are grateful to Xu Jin-Rong for sharing unpublished data on *CPK2*. Research in the Naqvi group is funded by the Temasek Life Sciences Laboratory (Singapore) and The National Research Foundation (Prime Minister’s Office, Grant Numbers NRF-CRP7-2010-02 and NRF-CRP16-2015-04), Singapore.

